# Liver-specific LXR inhibition represses reverse cholesterol transport in cholesterol-fed mice

**DOI:** 10.1101/2023.02.06.527401

**Authors:** Takafumi Nishida, Makoto Ayaori, Junko Arakawa, Yumiko Suenaga, Kazusa Shiotani, Harumi Uto-Kondo, Tomohiro Komatsu, Kazuhiro Nakaya, Yasuhiro Endo, Makoto Sasaki, Katsunori Ikewaki

## Abstract

**Objective:** High density lipoprotein (HDL) exerts an anti-atherosclerotic effect via reverse cholesterol transport (RCT). Several phases of RCT are transcriptionally controlled by Liver X receptors (LXRs). Although macrophage LXRs reportedly promote RCT, it is still uncertain whether hepatic LXRs affect RCT *in vivo*.

**Approach and Results:** To address this question, we induced hepatic overexpression of sulfotransferase family cytosolic 2B member 1 (Sult2b1) in mice. Sult2b1 facilitates generation of sulfated cholesterol, resulting in reduced production of LXR ligands (oxysterols), which impairs LXR signaling. Adenoviral vectors expressing Sult2b1 (Ad-Sult2b1) or luciferase were intravenously injected into mice under a normal or high-cholesterol diet. Hepatic Sult2b1 overexpression resulted in reduced expression of LXR-target genes - ATP-binding cassette transporter G5/G8, cholesterol 7α hydroxylase and LXRα itself - respectively reducing or increasing cholesterol levels in HDL and apolipoprotein B–containing lipoproteins (apoB-L). A macrophage RCT assay revealed that Sult2b1 overexpression inhibited fecal excretion of macrophage-derived ^3^H-cholesterol only under a high-cholesterol diet. In a HDL kinetic study, Ad-Sult2b1 promoted catabolism/hepatic uptake of HDL-derived cholesterol, thereby reducing fecal excretion. We next performed an *in vitro* lipoprotein production assay which revealed a Sult2b1-mediated reduction/increase in HDL or apoB-L secretion from hepatocytes, respectively. Finally, in LXRα/β double knockout mice, hepatic Sult2b1 overexpression increased apoB-L levels, but there were no differences in HDL levels or RCT compared to the control, indicating that Sult2b1-mediated effects on HDL/RCT and apoB-L were distinct: the former was LXR-dependent, but not the latter.

**Conclusions:** Hepatic LXR inhibition negatively regulates circulating HDL levels and RCT by reducing LXR-target gene expression.

**Graphic Abstract:** 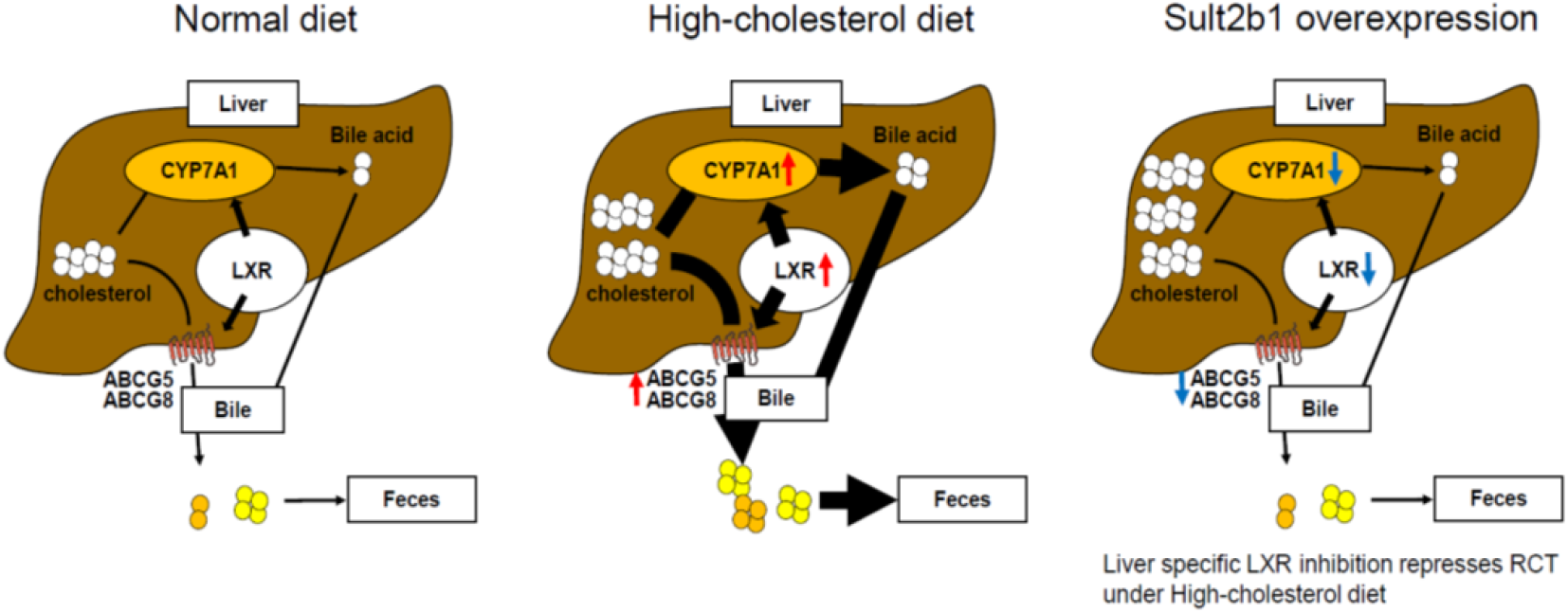

---

High density lipoprotein (HDL) has atheroprotective properties including reverse cholesterol transport (RCT).^1^ HDL promotes efflux of excess cholesterol from macrophage-derived foam cells and transports it back to the liver for excretion from bile to feces.^2^ Several steps of RCT are transcriptionally regulated by liver X receptors (nuclear receptors, LXRs) in macrophages, the liver, and intestine. LXRs are activated by oxysterols as natural ligands and induce LXR-target genes such as ATP-binding cassette transporter (ABC) A1 (ABCA1), ABCG1, ABCG5, ABCG8, and cholesterol 7α hydroxylase (CYP7A1). Pharmacological activation of LXRs using their agonists reportedly inhibits the development of atherosclerosis via activation of RCT in animal models.^3–6^ LXRβ is expressed ubiquitously whereas, LXRα is predominantly expressed in tissues, such as those of the liver, intestine and macrophages, which play important roles in cholesterol homeostasis. In whole body LXRα/β double-knockout mice (DKO), reduced HDL cholesterol (HDL-C) levels, attenuated RCT, and enhanced development of atherosclerosis were observed.^7, 8^ Further, accumulating evidence suggests beneficial effects of tissue-specific LXR expression on atherogenesis since macrophage or liver-specific LXR deletion resulted in accelerated development of atherosclerosis in mice. ^5, 8–10^

Macrophage LXRs reportedly contribute to anti-atherogenesis by reducing RCT^11^ although physiological roles of liver LXRs in RCT remain unclear. Hepatic LXRα deletion abolished LXR agonist-induced promotion of RCT; however, there was no difference in RCT between LXRα knockout and wild type mice under physiological conditions (without LXR agonist administration). ^10, 11^ Moreover, it remains uncertain how hepatic LXRβ contributes to RCT *in vivo*.

In order to extend our knowledge of the effects of hepatic LXRs on RCT, we induced hepatic overexpression of sulfotransferase family cytosolic 2B member 1 (Sult2b1) in mice to inhibit hepatic LXR pathways. Sult2b1 facilitates the generation of sulfated cholesterol by adding a sulfate group to cholesterol side chains, resulting in reduced oxysterol production, which, in turn, inhibits LXRα/β signaling.^12^ Here we demonstrate that hepatic LXR inhibition by Sult2b1 overexpression reduced circulating HDL levels and attenuated RCT but only under a high cholesterol (HC) diet.

## Materials and Methods

### Animals and Diet

C57BL/6J mice were obtained from Clea Japan (Tokyo, Japan) and fed a normal chow (NC) diet or a HC diet supplemented with 0.5% cholesterol (Oriental Yeast, Tokyo, Japan). Experiments were performed using mice aged 6 to 8 weeks. Mice were handled according the guidelines of National Defense Medical College Institutional Animal Care and Use Committee. In accordance with a previous study by Chen et al,^12^ we used a 0.5% HC diet for three weeks to activate LXR signaling pathways. We also used DKO, which were kindly provided by Dr Makoto Makishima of Nihon University.

### Generation of Recombinant Adenoviruses Encoding mouse Sult2b1

A recombinant adenovirus expressing mouse Sult2b1 (Ad-Sult2b1) was produced using the ViraPower Adenoviral Expression System (Invitrogen, Carlsbad, CA), according to the manufacturer’s instructions. Briefly, to generate an entry clone of the Gateway system (Invitrogen), cloning of the open reading frame into a pENTR/D-TOPO vector (Invitrogen) was carried out using first strand complementary DNA (cDNA) derived from a cDNA clone (Thermo Fisher Scientific, Waltham, MA) containing an open reading frame of Sult2b1 as a template, Platinum Pfx DNA polymerase (Invitrogen), and the following specific primers: forward: 5’ -CAC CAT GGA CGG GCC GCA GCC-3’; reverse: 5’ -TTA TTG TGA GGA TCC TGG GTT GGG-3’.

An expression clone for the adenoviral vector was then generated by performing a LR recombination reaction between the entry clone and pAd/CMV/V5-DEST (Invitrogen) according to the manufacturer’s protocol. The recombinant adenoviral plasmid was purified, and then transfected into 293A cells (Riken cell bank, Ibaraki, Japan). After a sufficient cytopathic effect was observed in 293A cells, the adenovirus was purified using the Fast Trap Adenovirus Purification and Concentration Kit (Merck KGaA, Darmstadt, Germany). An adenoviral vector expressing luciferase (Ad-Luc) was kindly donated by Dr. Santamarina-Fojo S of National Institute of Health and used as a control. The adenovirus titer in plaque-forming units (PFU) was determined using a Adeno-X Rapid Titer kit (Clontech, Palo Alto, CA).

### Plasma Lipid and Lipoprotein Analysis

Blood sampling was performed on the indicated days after injection of Ad-Sult2b1 or -Luc (7.5×10^8^ PFU) into the mice. Plasma total cholesterol (TC) levels were measured using commercially available assay kits (Wako Pure Chemical Industries, Ltd., Osaka, Japan). For lipoprotein fractionation analysis, equal volumes of plasma samples were pooled from each group of mice (total volume of 500 μL). Lipoproteins were fractionated using a Superose 6 10/300 GL fast protein liquid chromatography (FPLC) column (GE healthcare, UK). Fractions (500 μL) were collected and used for lipid measurement.

### Western Blot Analyses

On the indicated days after injection of Ad-Sult2b1 or -Luc, livers and plasma were obtained. Protein extracts were prepared using T-PER (Pierce Chemical Co., IL) in the presence of protease inhibitors (Roche Applied Science, Barcelona, Spain), and subjected to Western blot analyses with anti-rat anti-ABCA1 antiserum (kindly donated by Dr. S. Yokoyama of Nagoya City University, Japan),^13^ anti-rabbit anti-scavenger receptor class B type I (SR-BI) and anti-rabbit anti-LCAT (Novus Biologicals, CO), anti-mouse β-actin, anti-rabbit anti-ABCG8, anti CYP7A1, anti-low density lipoprotein (LDL) receptor (LDLR) and anti-LXRα (Abcam, Cambridge, UK), anti-rabbit anti-sterol regulatory element-binding protein-2 (SREBP2, Cayman chemical, MI), anti PCSK9 (Cayman chemical), anti-rabbit anti-ABCG5, anti-mouse anti-apolipoprotein (apo) A-I (Santa Cruz, CA), and anti-rabbit anti-endothelial lipase (EL, National Institute of Health, MD) specific antibodies. The proteins were visualized by a chemiluminescence method (Amersham Imager 600, GE healthcare Life Sciences, Buckinghamshire, UK). Protein signals were quantified using ImageJ software (National Institute of Health, MD), and expressed as the ratio of individual protein signals to β-actin signals.

### Real-time Quantitative RT-PCR

Total RNA was extracted from the livers and small intestines, and first-strand cDNA was synthesized from the total RNA (500 ng) by placing in a Reverse Transcription Reagent (Applied Biosystems, Foster City, CA). Quantitative PCR was performed with a Perkin–Elmer 7900 PCR machine, TaqMan PCR master mix and FAM-labeled TaqMan probes (Assays-on-Demand, Applied Biosystems) for mouse LXRα, ABCA1, ABCG5, ABCG8, CYP7A1, PCSK9, LDLR, SREBP2, SR-BI, and 18S. Expression data were normalized for 18S levels.

### *In Vivo* Macrophage RCT Studies

RAW264.7 cells were grown in RPMI1640 supplemented with 10% fetal bovine serum (FBS), and then radiolabeled with 5 μCi/mL ^3^H-cholesterol and cholesterol enriched with 10 μg/mL of acetylated LDL (AcLDL, Invitrogen, MA) for 48 hr. These foam cells were washed, equilibrated, detached with cell scrapers, resuspended in RPMI 1640, and pooled before being injected into mice.

The mice were caged individually with unlimited access to food and water. Seven days after intravenous injection of adenoviral vectors, ^3^H-cholesterol-labeled and AcLDL-loaded RAW264.7 cells (typically 5.0 × 10^6^ cells containing 7.5 × 10^6^ counts per minute [dpm] in 0.5 mL RPMI 1640) were injected intraperitoneally as described previously.^14^ Blood was obtained at 24 and 48 hr, and plasma was subjected to liquid scintillation counting (LSC). Feces were collected continuously from 0 to 48 hr and stored at 4°C before being counted. At 48 hr after injection, mice were exsanguinated to examine livers, intestines, and bile.

Liver lipids were extracted according to the procedure of Bligh and Dyer.^15^ Briefly, a 50 mg piece of tissue was homogenized in water, and lipids were extracted with a 2:1 (vol/vol) mixture of chloroform/methanol. The lipid layer was collected, evaporated, resuspended in a 3:2 (vol/vol) mixture of hexane/isopropanol, and counted by LSC. Bile was directly counted by LSC. Total feces collected from 0 to 48 hr were weighed and soaked in Millipore water (1 mL water per 100 mg feces) overnight at 4 °C. An equal volume of ethanol was added the next day, and samples were homogenized, 200 μL of which were counted by LSC. Results were expressed as percentage of dpm injected. To extract ^3^H-neutral sterols and ^3^H-bile acid fractions, 2 mL of the homogenized samples was combined with 2 mL ethanol, and 400 μL sodium hydroxide. The samples were saponified at 95°C for 2 hr, cooled to room temperature, and then ^3^H-cholesterol was extracted 3 times with 9 mL hexane. The extracts were pooled, evaporated, resuspended in toluene, and then counted by LSC. To extract ^3^H-bile acids, the remaining aqueous portion of the feces was acidified with concentrated hydrogen chloride and then extracted 3 times with 9 mL ethyl acetate. The extracts were pooled, evaporated, resuspended in ethyl acetate, and counted by LSC.

### HDL Metabolic Studies

Human HDL was isolated from pooled human plasma by sequential ultracentrifugation (density 1.063 < density 1.21 g/mL). Dialyzed human HDL was labeled with ^3^H-cholesteryl oleate (CEs). Fifty μCi of CEs in toluene were dried down under nitrogen. Ethanol was then added, and the solution taken up by a pipette. This was added to the HDL solution (1 mL of dialyzed HDL containing of 10 mg of protein) over a period of 5 min while gently shaking with short interruptions for a brief vortex. The HDL solution was incubated for 24 hr at 37 °C. The HDL from the solution was re-isolated by ultracentrifugation (40,000 rpm, 48 hr) at the original density and the ^3^H-HDL was dialyzed overnight against phosphate-buffered saline (PBS) containing 0.01% EDTA. Finally, the ^3^H-HDL was filter-sterilized and stored at 4 °C until injection.

For the metabolic study, 1 × 10^6^ dpm of HDL labeled with ^3^H-CEs were injected intravenously via tail veins into C57BL/6J mice at day 7 after Ad-Sult2b1 or Ad-Luc injection. Blood was collected from the tail vein at 2 min, 30 min, and at 1, 3, 6, 9, 24 and 48 hr. Feces were collected continuously up to 48 hr. Aliquots of 10 μl of plasma were counted for each time point by LSC. After correction with blank values, plasma decay curves for the tracer were normalized to the plasma counts at the initial 2 min time point after tracer injection. The fractional catabolic rates (FCR) were determined using the SAAMII program.^16^ After the 48 hr blood samples were collected, livers and bile were harvested. Results were expressed as a percentage of the injected dose.

### Lipoprotein Production Assay in Cultured Hepatocytes

Hepa 1-6 cells (ATCC, Manassas, VA) were maintained in Dulbecco’s modified Eagle’s medium (DMEM) containing 10% fetal bovine serum. They were seeded on 6-well plates to be more than 90% confluent. After preparation, the cells were treated with 30MOI of Ad-Sult2b1 or –Luc for 24 hr (each 3 wells), washed with PBS three times, and then incubated in DMEM containing ^3^H-cholesterol (5.0 μCi/mL) and 5% bovine serum albumin (BSA). After 24 hr, the supernatants were removed. The cells were washed with PBS three times and incubated in DMEM containing 5% BSA. After 24 hr, the supernatants were harvested, centrifuged, and then concentrated using an Amicon Ultra-15 filter unit (Merck KGaA, Darmstadt, Germany). After lipoprotein fractionation of the concentrated media using FPLC, the ^3^H-tracer activity in each fraction was measured.

### Statistical Analysis

All the statistical analyses were conducted using the 2-tailed Student’s t-test with GraphPad Prism Software. Results are presented as mean ± SD. p values of less than 0.05 were considered to be statistically significant.

## Results

### Intravenous Injection of Adenoviral Vector Encoding Sult2b1 Resulted in Liver-Specific Overexpression of Sult2b1 and Suppressed ABCG5/G8 and CYP7A1 expression

First, we intravenously injected adenoviral vectors encoding mouse Sult2b1 (Ad-Sult2b1) and luciferase (Ad-Luc), as a control, into C57BL/6J mice and determined Sult2b1 mRNA levels in the liver and small intestine 9 days after the injection. Transduction with Ad-Sult2b1 resulted in Sult2b1 mRNA overexpression in the liver, but not in the small intestine (Supplemental Figure I), indicating that intravenous injection of adenoviruses achieved liver-specific overexpression of Sult2b1. We also measured mRNA levels of hepatic molecules involved in cholesterol metabolism. Under a NC diet, Sult2b1 overexpression did not affect expression of LXR target genes such as ABCG5, ABCG8, and CYP7A1 (Supplemental Figure II). HC resulted in enhanced ABCG5 levels, which were canceled by Sult2b1 overexpression, while Ad-Sult2b1 transduction reduced ABCG8 and CYP7A1 levels only under HC, supporting previous observations.^12^ Sult2b1 overexpression repressed LXR signaling pathways in the liver. In contrast, expressions of other LXR-target genes, ABCA1 and LXRα itself, were increased by Ad-Sult2b1. Sterol regulatory element-binding protein-2 (SREBP2) and expression of its target genes, low density lipoprotein (LDL) receptor (LDLR) and proprotein convertase subtilisin/kexin type 9 (PCSK9), were reduced under HC as expected, but not changed by Sult2b1 overexpression.

Western blot analyses revealed that Ad-Sult2b1 treatment reduced hepatic LXRα, ABCG5/G8 and CYP7A1 expressions in mice fed with either the NC or HC diet (Figure 1A). Differing from its mRNA levels, SREBP2 protein expression was dramatically reduced by Ad-Sult2b1. PCSK9 protein levels, however, were not affected by diet or Ad-Sult2b1. Contrary to the changes in its mRNA levels, LDLR protein levels were unexpectedly increased by Ad-Sult2b1 treatment. There was no obvious change in ABCA1 and scavenger receptor class B type I (SR-BI) expression with or without HC diet or Sult2b1 overexpression. Although serum EL expression was increased by Ad-Sult2b1 treatment under the NC diet, there was no difference in this respect between Ad-Sult2b1 and Ad-Luc under the HC diet.

**Figure 1.**
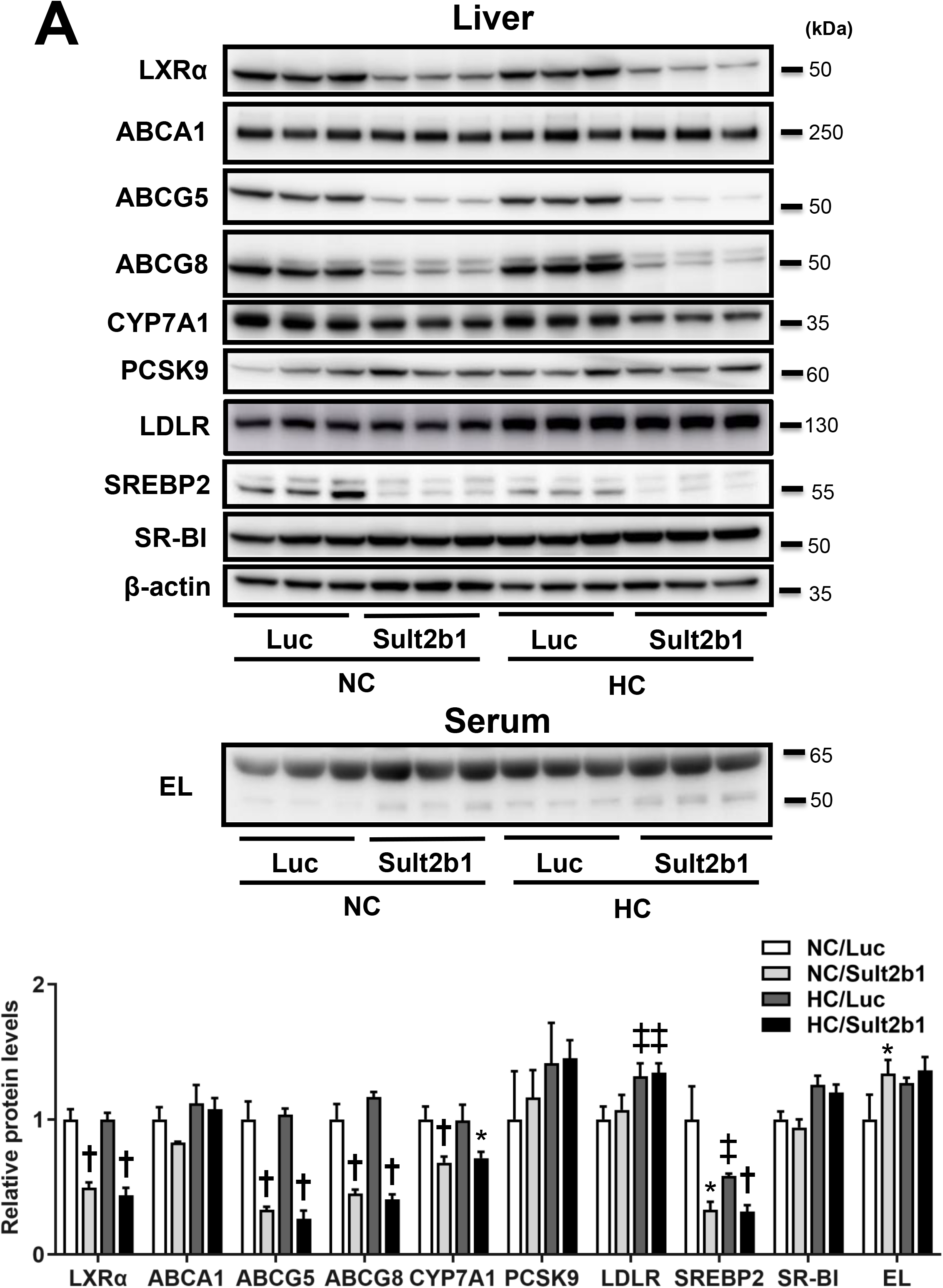
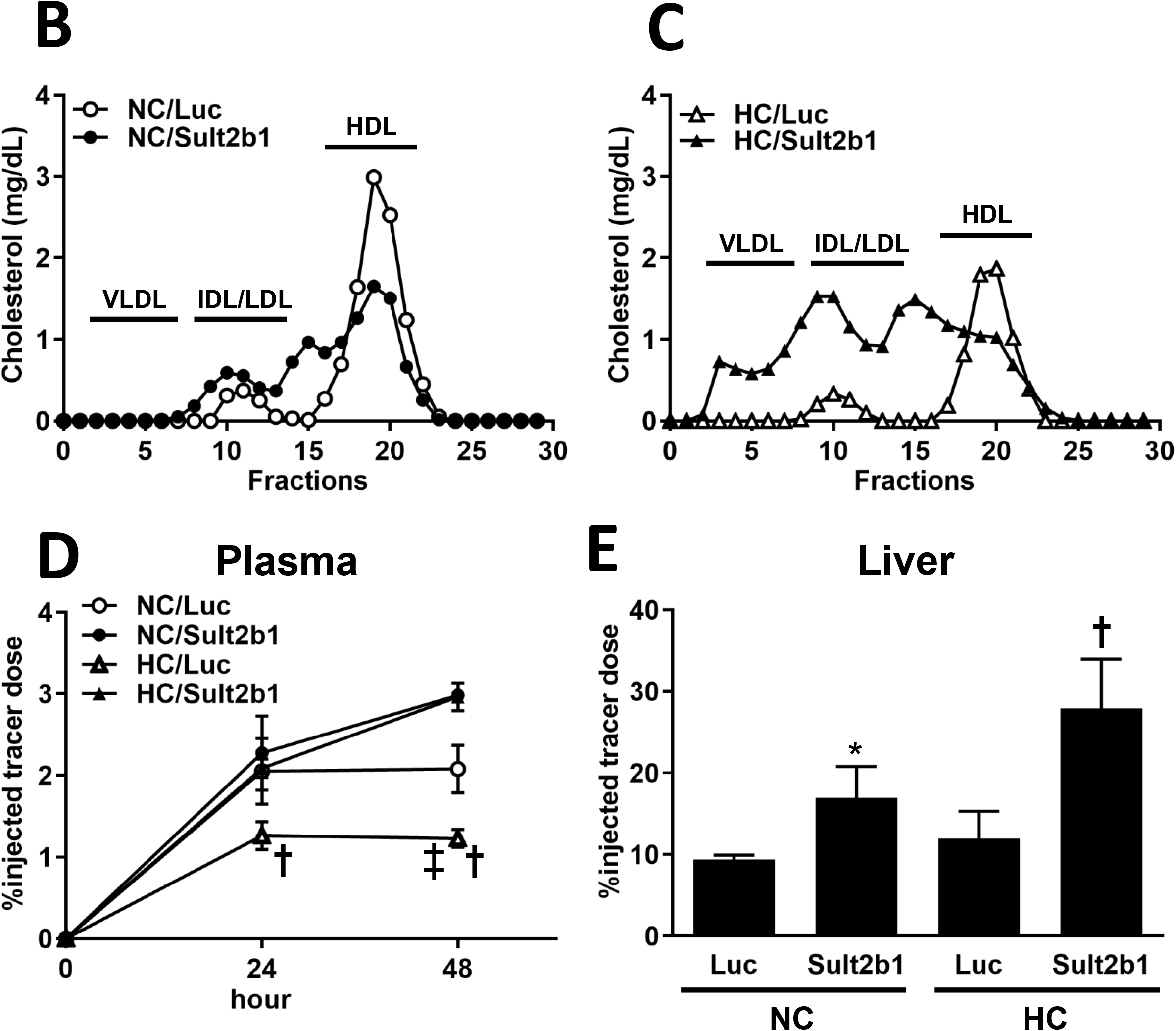
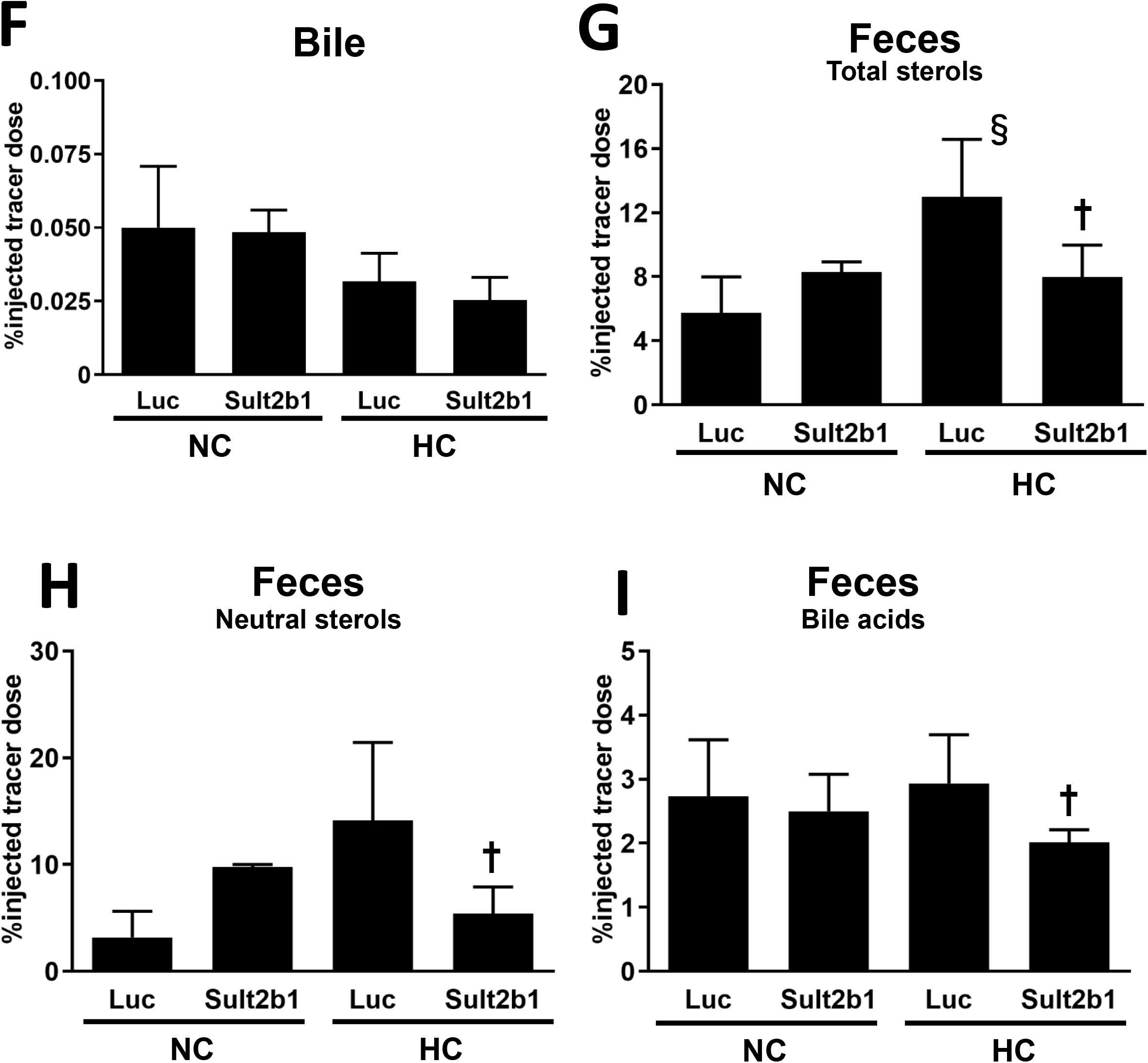
Hepatic Sult2b1 overexpression reduced macrophage RCT only under high cholesterol (HC) chow. A, Nine days after intravenous injection of Ad-Sult2b1 or -Luc into C57BL/6J mice under normal chow (NC) or HC chow, liver samples were obtained and subjected to Western blot analyses as described in Materials and Methods. B-C, Nine days after intravenous injection of Ad-Sult2b1 or -Luc into C57BL/6J mice under NC and HC chow, plasma was obtained and separated into lipoprotein fractions by fast protein liquid chromatography (FPLC). Cholesterol levels in each fraction were determined as described in Materials and Methods. D-G, Seven days after intravenous injection of Ad-Sult2b1 or -Luc into C57BL/6J mice under NC and HC chow, ^3^H-cholesterol-labeled RAW cells were intraperitoneally injected. At the indicated hours after injection, plasma was obtained and subjected to ^3^H-tracer analysis (D). Forty-eight hr after injection, the liver (E) and bile (F) were isolated, and then subjected to ^3^H-tracer analysis. Feces (G, H, I) collected continuously from 0 to 48 hr were subjected to ^3^H-counting. Neutral sterols and bile acids were separated as described in Materials and Methods. Data are expressed as percent counts relative to total injected tracer (means ± SD, n=8 for each group). * p<0.05, † p<0.01 vs. Luc. ‡ p<0.05, § p<0.01 vs. NC.

### Hepatic Sult2b1 Overexpression Reduced HDL-C and Increased ApoB-containing Lipoprotein Levels

As shown in Figure 1B and 1C, FPLC analysis of plasma obtained 9 days after virus injection revealed that HDL-C levels were decreased under HC, as compared to NC, and further reduction was observed with hepatic Sult2b1 overexpression. Cholesterol levels in apolipoprotein (apo) B-containing lipoproteins were unchanged by hepatic Sult2b1 overexpression under NC. In contrast, under HC they were dramatically increased by Ad-Sult2b1.

### Hepatic LXR Inhibition by Sult2b1 Repressed RCT Only under HC Diet

The effects of hepatic LXR inhibition induced by Sult2b1 overexpression on RCT were assessed by performing *in vivo* macrophage RCT studies. Briefly, after intraperitoneally injecting ^3^H-cholesterol labeled macrophages, radioactivity was measured in plasma, liver, bile, and feces. As shown in Figure 1D, HC markedly reduced the appearance of macrophage-derived ^3^H-cholesterol in the circulation. The reduction was cancelled by Sult2b1 overexpression, and was not observed under NC. To assess the distribution of macrophage-derived ^3^H-cholesterol, we measured radiotracer activities in lipoprotein fractions separated by FPLC. This showed that distribution of macrophage-derived ^3^H-cholesterol was mostly identical to that of non-radiolabeled cholesterol (Figure 1B & C, Supplemental Figure III), suggesting that reduced ^3^H-cholesterol levels in plasma under HC mirrored the changes in HDL. In contrast, Ad-Sult2b1 restored tracer levels, and this was due to an increase in apoB-containing lipoproteins. Regarding changes in the liver and bile, Sult2b1 overexpression increased liver uptake of ^3^H-cholesterol, which was further enhanced by HC (Figure 1E), while Ad-Sult2b1 did not affect radioactivity in the bile under either diet (Figure 1F). Fecal excretion of ^3^H-tracer, the determinant of *in vivo* RCT, was increased under HC as compared to NC, and the increase was cancelled by Ad-Sult2b1 (Figure 1G: 5.8±2.2% versus 13.0±3.6% versus 8.0±2.0%; NC/Luc versus HC/Luc versus HC/Sult2b1, respectively). On the other hand, fecal excretion of ^3^H-tracer did not differ between the viruses under NC. We also determined fecal tracer levels in the neutral sterol and bile acid subfractions, revealing that the reduction in neutral sterols was more remarkable than that in bile acids (Figure 1H & I).

### Hepatic Sult2b1 Overexpression Promoted Catabolism of HDL and Reduced Fecal Excretion of HDL-Derived Cholesterol

To further investigate the underlying mechanisms for reduced HDL-C levels and RCT in Ad-Sult2b1-treated mice, we performed an HDL turnover study using HDL labelled with ^3^H-cholesteryl oleate (CEs), which is metabolized to free cholesterol and bile acids in the liver. Figure 2A shows that HDL-CEs was catabolized much faster in mice fed with the HC diet than those under NC, and in Ad-Sult2b1-treated mice, HDL catabolism was further accelerated as compared to Ad-Luc, with fractional catabolic rates (FCRs) of 0.10±0.01, 0.15±0.02, and 0.19±0.02 pools/day, respectively (Figure 2B). These results suggest that reduction in HDL-C levels was attributable to increased catabolism of HDL both under HC and hepatic Sult2b1 overexpression. Regarding distribution of HDL-derived ^3^H-CEs in the liver and bile, findings were similar to those for macrophage-derived ^3^H-cholesterol; i.e., hepatic Sult2b1 overexpression caused increased tracer levels in the liver, but not in bile (Figure 2C & D). Finally, the fecal excretion of HDL-derived HDL-CEs increased under HC as compared to NC and the increase was cancelled in Sult2b1-overexpressing mice (Figure 2E: 8.0±1.9, 14.7±2.8, 9.9±2.5%; NC/Luc, HC/Luc, HC/Sult2b1, respectively). Taken together with observations in the experiments on distribution of macrophage-derived ^3^H-cholesterol (Figure 1), these results suggest that the first step of RCT, from macrophages to HDL, did not contribute to the changes in RCT under HC and Sult2b1 overexpression.

**Figure 2.**
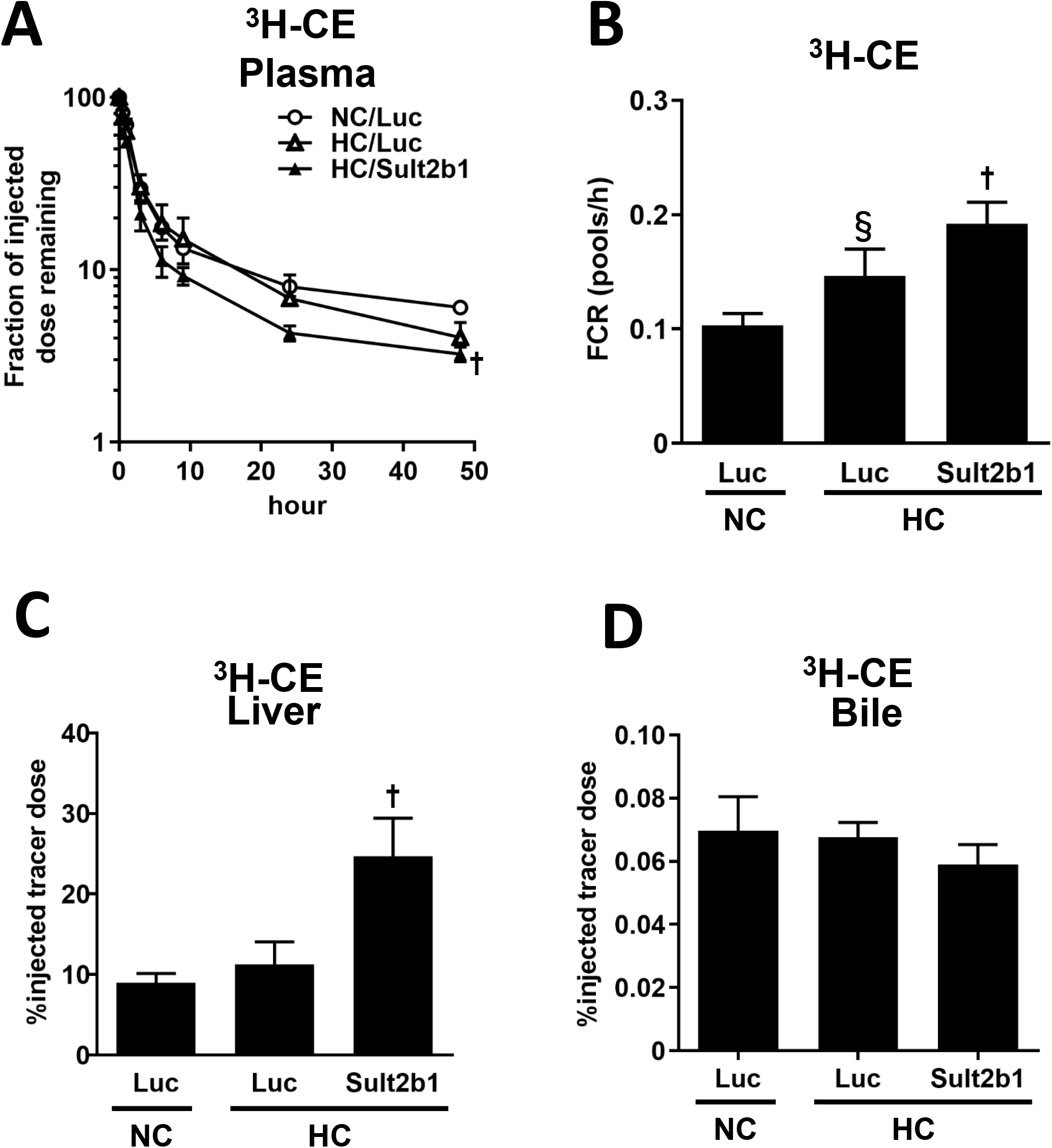
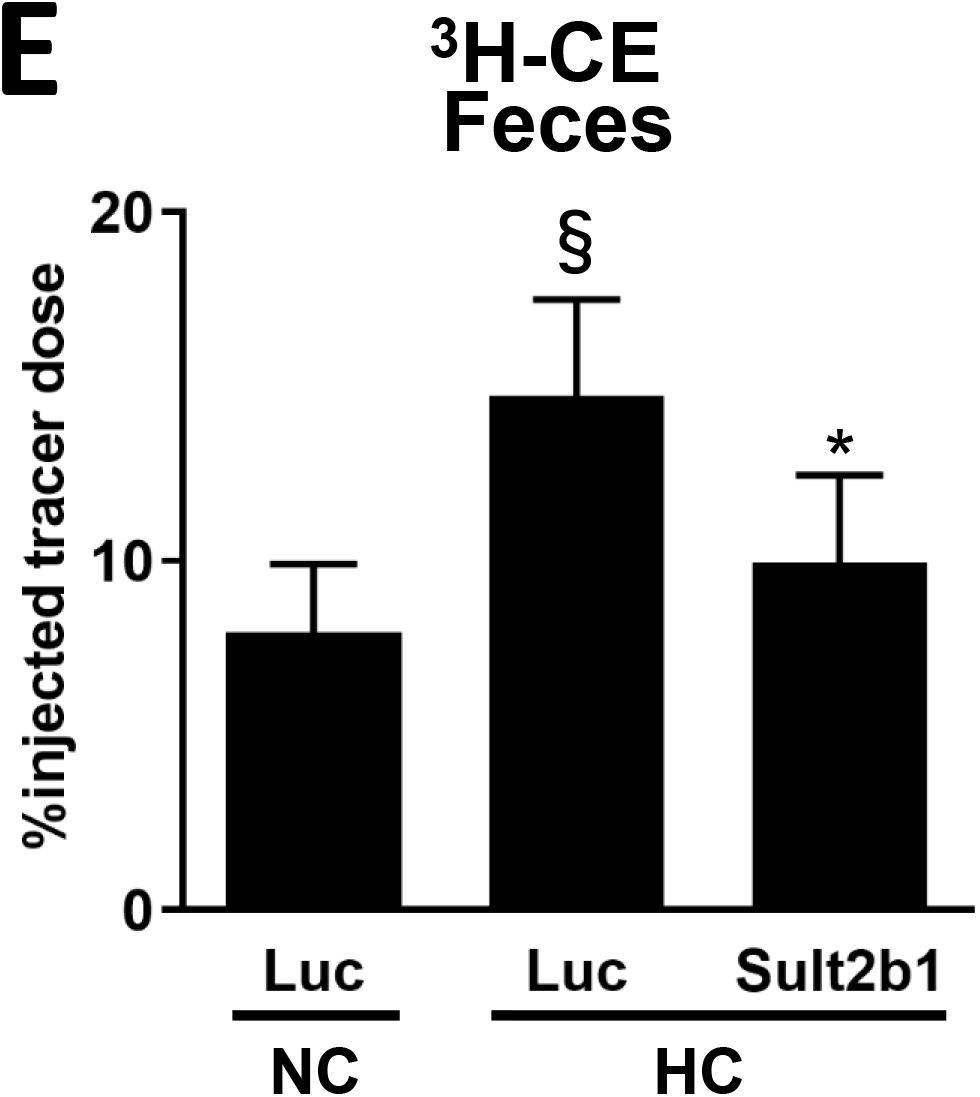
Hepatic overexpression of Sult2b1 accelerates HDL catabolism, increases hepatic uptake of HDL-cholesteryl-oleate, and reduces fecal excretion under HC chow. A, Seven days after intravenous injection of Ad-Sult2b1 or -Luc, the mice were intravenously injected with ^3^H-Cholesteryl oleate (CEs)-labeled HDL. Plasma was obtained after the indicated times, and then subjected to ^3^H-tracer analysis. Plasma decay curves were normalized to radioactivity at the initial 2-min time point after tracer injection. Data are expressed as percent counts relative to total injected tracer, mean ± SD. B, Fractional catabolic rates (FCR) were determined from the area under the plasma disappearance curves fitted to a bi-compartmental model using the SAAM II program. C-E, Forty-eight hr after injection with ^3^H-CEs-HDL, livers (C) and bile (D) were isolated and then subjected to ^3^H-tracer analysis. Feces collected continuously from 0 to 48 hr were also subjected to ^3^H-counting (E). Values are expressed as percentage of the total ^3^H-CEs-HDL injected (means ± SD, n=6 for each group). * p<0.05, † p<0.01 vs. Luc. ‡ p<0.05, § p<0.01 vs. NC.

### Sult2b1 Overexpression Respectively Increased/Decreased VLDL/HDL Production by Cultured Hepatocytes

We next investigated whether LXR pathway inhibition affected hepatic lipoprotein production using cultured hepatocytes preincubated with ^3^H-cholesterol. As shown in Figure 3, VLDL and HDL production was respectively increased and decreased in Sult2b1-overexpressing hepatocytes as compared to the control, suggesting that increased and reduced production of VLDL and HDL were respectively attributable to increased and reduced circulating VLDL and HDL levels in Ad-Sult2b1-treated mice.

**Figure 3.**
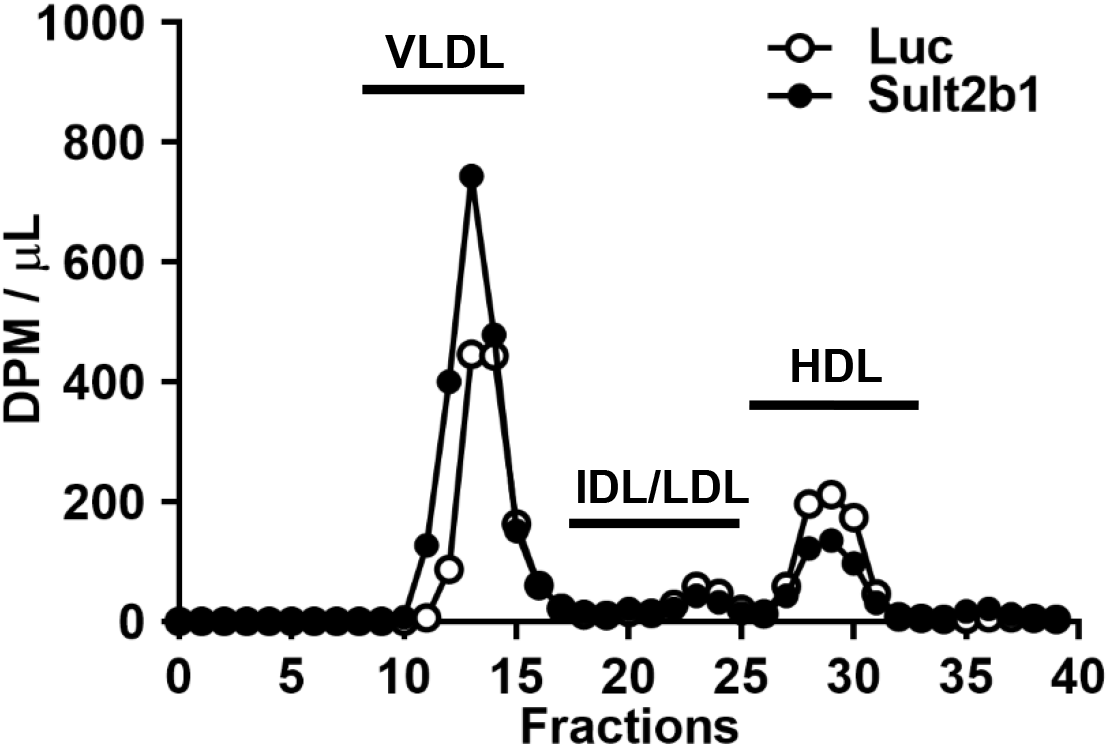
Sult2b1 overexpression respectively increases/reduces VLDL/HDL production by cultured hepatocytes. Hepa 1-6 cells were treated with 30MOI of Ad-Sult2b1 or –Luc for 24 hours, washed, and then incubated with ^3^H-cholesterol containing media. After 24 hr, the supernatants were centrifuged, concentrated, and applied to fast protein liquid chromatography (FPLC) to separate lipoprotein fractions, which were subjected to ^3^H-tracer analysis as described in Materials and Methods. To eliminate the possibility that the labeling efficiency of ^3^H-cholesterol differed, we confirmed that there was no difference in trace levels between the adenoviruses 24 hr after labeling.

### Circulating HDL-C Levels and RCT Were Reduced in DKO

To eliminate the possibility that hepatic Sult2b1 overexpression affects lipoprotein profiles via LXR-independent pathways, we performed the experiments under HC using DKO. Supplemental Figure IV shows that hepatic mRNA levels of the LXR target genes ABCG5/G8, and CYP7A1 were lower in DKO than in the wild-type. Sult2b1 overexpression, however, did not affect mRNA expression of these molecules, indicating that the effects of Sult2b1 overexpression in this respect were mediated via LXR-dependent pathways. Although Ad-Sult2b1 treatment did not affect PCSK9 mRNA levels (Supplemental Figure II), whole body LXRα/β deletion resulted in reduced PCSK9 expression, which was not affected by Ad-Sult2b1. There were no differences in ABCA1, SREBP2, LDLR or SR-BI mRNA levels between wild-type mice and DKO.

First, we compared protein expressions, lipid profiles and RCT between wild-type mice and DKO. Figure 4A shows that, under HC, ABCG5/G8 protein levels in DKO were lower as compared to those in the wild-type. In contrast, ABCA1, CYP7A1 and SREBP2 protein levels did not differ between wild-type mice and DKO. FPLC analysis revealed that whole body LXRα/β deletion reduced HDL-C levels but slightly increased apoB-containing lipoproteins as compared to wild-type mice (Figure 4B), findings consistent with those of previous studies.^11, 17, 18^ Similar to the findings in Figure 1D, LXR inhibition reduced appearance of macrophage-derived cholesterol in plasma (Figure 4C), in parallel with decreased circulating HDL-C levels. Concerning the distribution of ^3^H-cholesterol in the liver and bile, we obtained results distinct from those in Ad-Sult2b1 overexpressing mice. Namely, LXR deletion under HC reduced tracer levels in the bile and did not affect them in the liver (Figure 4D & E); in sharp contrast to Ad-Sult2b1 treatment, when there was an increase in the liver but no change in the bile (Figure 1E & F). Fecal excretion of macrophage-derived cholesterol was markedly reduced in DKO as compared to wild-type mice (Figure 4F).

**Figure 4.**
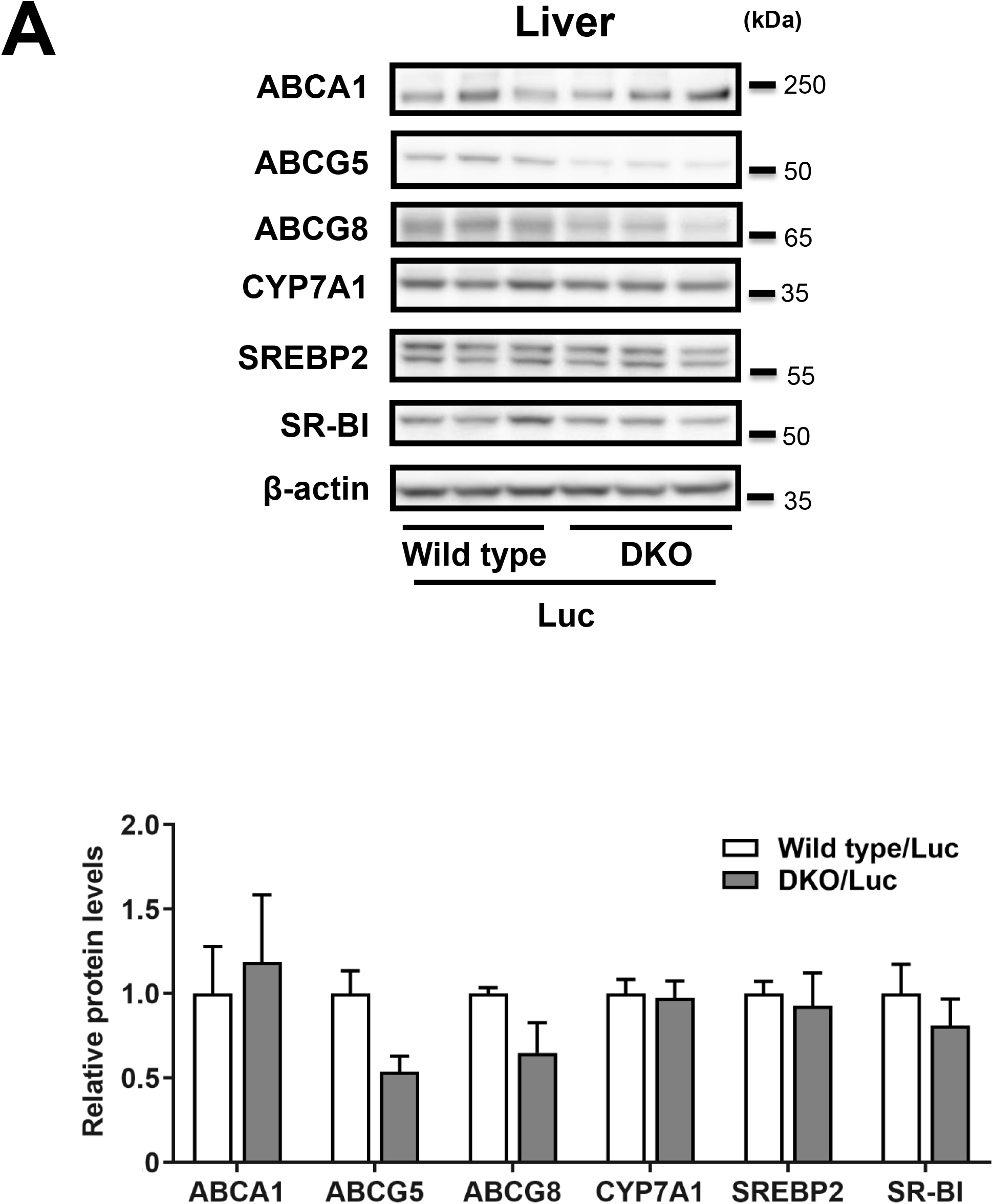
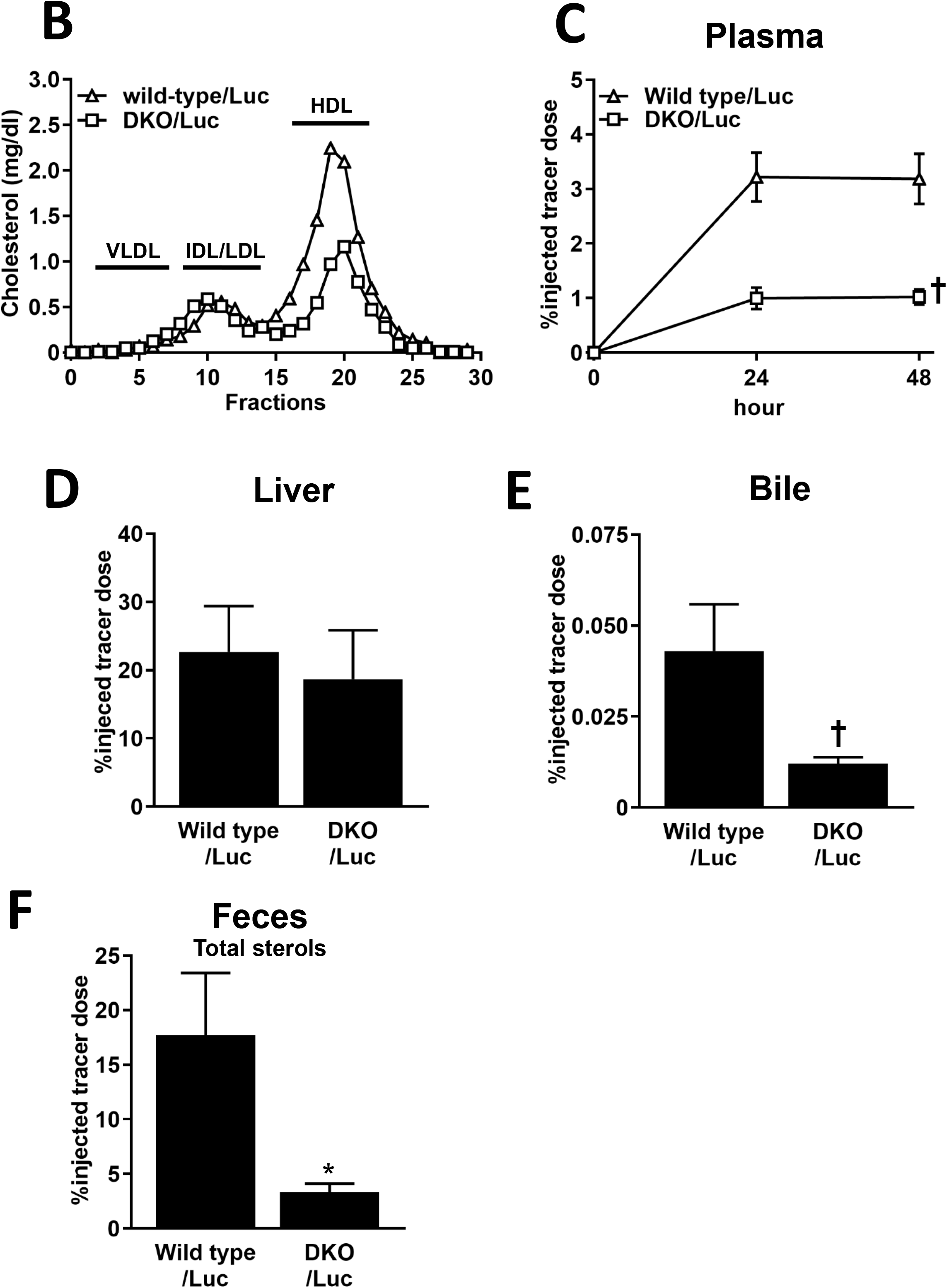
In LXRα/β double knockout mice (DKO), HDL-C levels and macrophage RCT were markedly reduced under HC chow. A, Nine days after intravenous injection of Ad-Luc into DKO or C57BL/6J mice (Wild type) under HC chow, liver samples were obtained and subjected to Western blot analyses as described in Materials and Methods. B, Nine days after intravenous injection of Ad-Luc into DKO or Wild type mice under HC chow, plasma was obtained and separated into lipoprotein fractions by FPLC. Cholesterol levels in each fraction were determined as described in Materials and Methods. C-F, Seven days after intravenous injection of Ad-Luc into DKO or the Wild type under HC chow, ^3^H-cholesterol-labeled RAW cells were intraperitoneally injected. At the indicated hours after injection, plasma was obtained and subjected to ^3^H-tracer analysis (C). Forty-eight hr after injection, the liver (D) and bile (E) were isolated, and then subjected to ^3^H-tracer analysis. Feces (F) collected continuously from 0 to 48 hr were subjected to ^3^H-counting. Data are expressed as percent counts relative to total injected tracer (means ± SD, n=8 for each group). * p<0.05, † p<0.01 vs. Wild type.

### Reduced HDL and Increased ApoB-containing Lipoprotein Levels Caused by Sult2b1 Overexpression Were Mediated by LXR-Dependent/-Independent Pathways, Respectively

Next, we investigated the effects of hepatic overexpression of Sult2b1 on lipid profiles and RCT in DKO under HC. In Western blot analysis, hepatic Sult2b1 overexpression did not affect ABCA1, ABCG5/G8, CYP7A1, SREBP2, or SR-BI protein levels as compared to the control (Figure 5A). FPLC analysis revealed that HDL-C levels were completely comparable between Ad-Sult2b1 and Ad-Luc –treated DKO while in sharp contrast, Ad-Sult2b1 transduction increased apoB-containing lipoprotein levels. Taken together with the observed FPLC pattern in Figure 4B, this indicates that increases in apoB-containing lipoprotein levels were partially mediated by LXR dependent pathways. On the other hand, Sult2b1 overexpression reduced HDL-C levels via LXR-dependent pathways.

**Figure 5.**
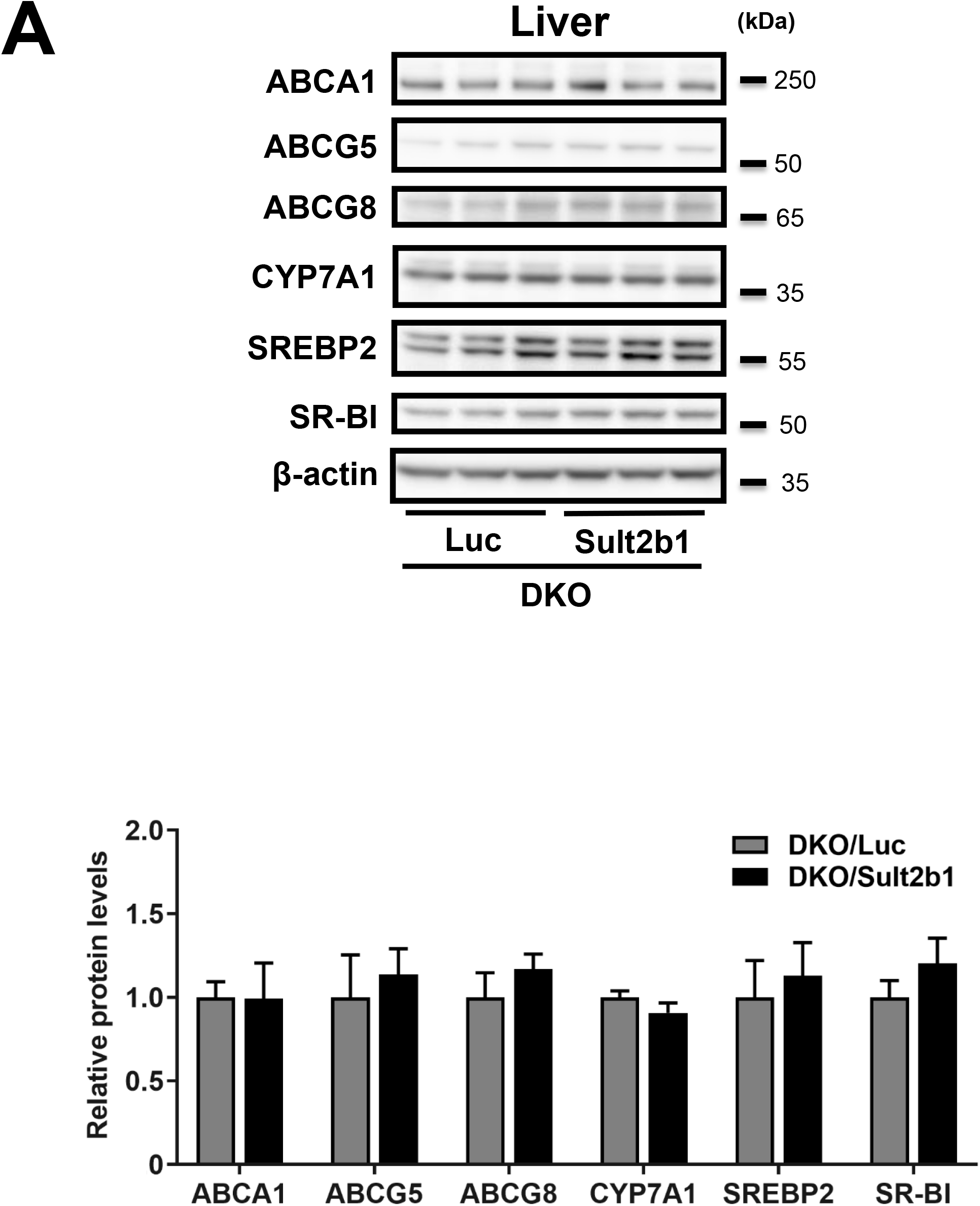
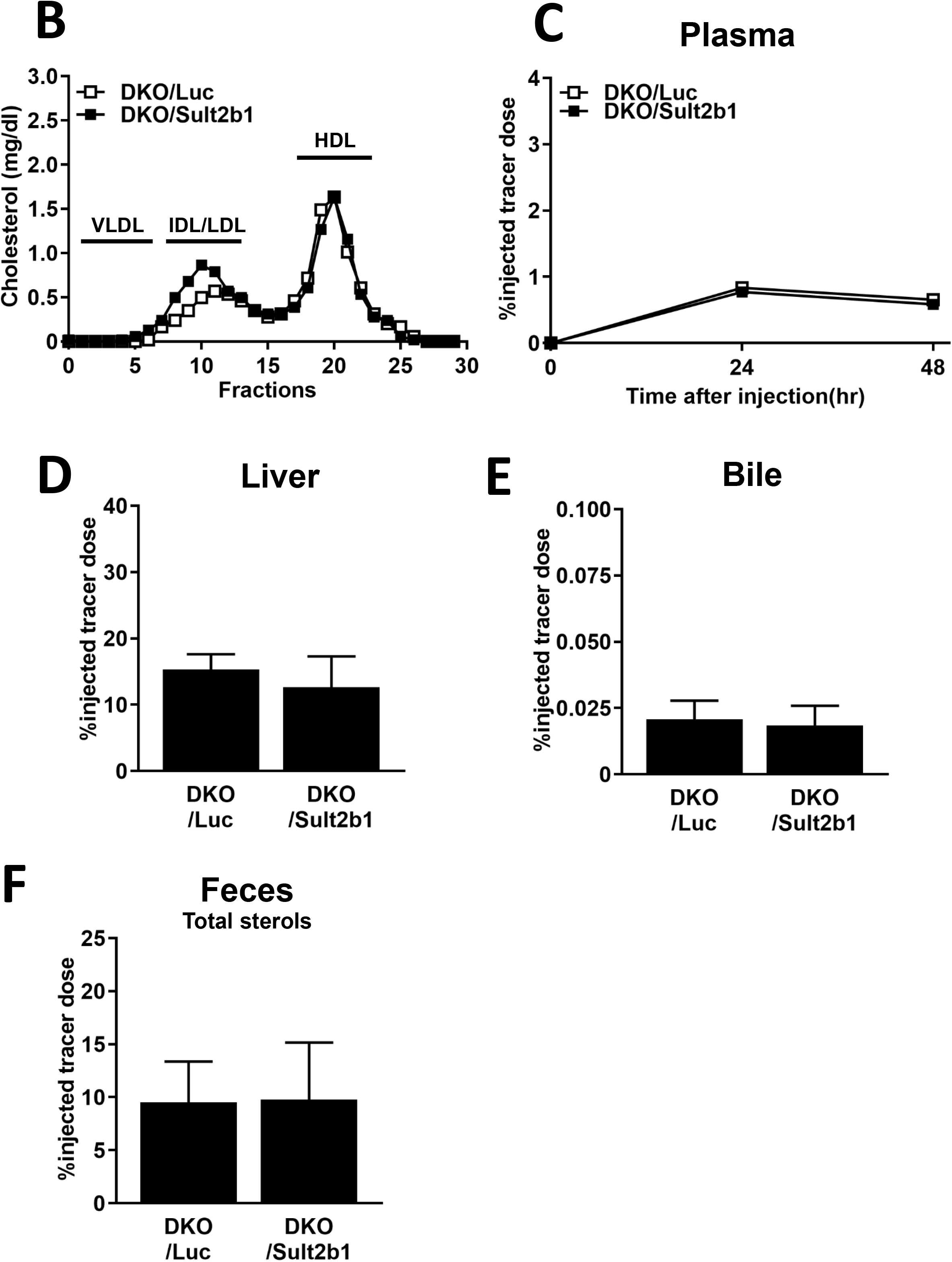
Hepatic Sult2b1 overexpression does not affect HDL-C levels or macrophage RCT in DKO under HC chow. A, Nine days after intravenous injection of Ad-Sult2b1 or -Luc into DKO under HC chow, liver samples were obtained and subjected to Western blot analyses as described in Materials and Methods. B, Nine days after intravenous injection of Ad-Sult2b1 or -Luc into DKO under HC chow, plasma was obtained and separated into lipoprotein fractions by FPLC. Cholesterol levels in each fraction were determined as described in Materials and Methods. C-F, Nine days after intravenous injection of Ad-Sult2b1 or -Luc into DKO under HC chow, ^3^H-cholesterol-labeled RAW cells were intraperitoneally injected. At the indicated hours after injection, plasma was obtained and subjected to ^3^H-tracer analysis (C). Forty-eight hours after injection, the liver (D) and bile (E) were isolated, and then subjected to ^3^H-tracer analysis. Feces (F) collected continuously from 0 to 48 hr were subjected to ^3^H-counting. Data are expressed as percent counts relative to total injected tracer (means ± SD, n=8 for each group).

### Hepatic Sult2b1 Overexpression Did Not Affect RCT in DKO

Finally, to investigate whether the effects of hepatic overexpression of Sult2b1 on RCT were dependent on LXRs, we performed an *in vivo* RCT assay using DKO. As shown in Figure 5C through 5F, Sult2b1 overexpression did not affect appearance of ^3^H-cholesterol in the plasma, liver, bile, or feces, thus indicating that Sult2b1 overexpression inhibited RCT in an LXR-dependent fashion.

## Discussion

The large burden of atherosclerotic diseases remains despite currently available optimum medical therapies. LDL has been the primary target in lipid management and HDL has been considered as the next target to further reduce the residual risk. However, therapeutic strategies targeting circulating HDL levels, especially cholesteryl ester transfer protein inhibitors,^19^ have failed to provide clinical benefits. In this context, one would expect that enhancing HDL functionalities, in particular RCT, rather than HDL quantity could be the next target for atherosclerosis.

Based on accumulating evidence of their favorable properties of enhancing cholesterol efflux from various tissues and raising HDL-C levels, LXRs have been emerging targets for atherosclerotic diseases. However, systemic administration of LXR agonists reportedly causes adverse effects - fatty liver and hypertriglyceridemia - by activating hepatic SREBP1c. For this reason, several researchers have made efforts to explore gene-selective^20^ or organ-specific^17^ LXR modulators, and have found these compounds to be promising. In the present study, we inhibited hepatic LXRs through Sult2b1 overexpression and observed reduced circulating HDL levels and attenuated RCT under a HC diet in an LXR-dependent fashion.

In accordance with a previous study,^12^ we established a mouse model of liver-specific LXR inhibition by means of liver-specific Sult2b1 overexpression (Supplemental Figure I), where LXR-target genes including ABCG5/8 and CYP7A1 were cancelled under a HC diet (Supplemental Figure II). We demonstrated that hepatic LXR inhibition resulted in lowered circulating HDL-C levels, supporting the findings of previous studies in mice with systemic deletion (Figure 4B) ^11, 18^ and liver-specific ^11^ deletion of LXRs. However, these studies failed to elucidate why LXR inhibition induces low HDL-C levels. In the present study, for the first time, we performed two experiments to investigate whether hepatic Sult2b1 overexpression affects HDL production and/or catabolism. We demonstrated that both enhanced catabolism (Figure 2A) and reduced production (Figure 3) were responsible for reduced circulating HDL levels. However, it remains unclear which molecules affect these phenomena since we did not observe reasonable changes in the protein expressions of ABCA1 (associated with HDL production), or SR-BI/EL (HDL catabolism).^21^ Further studies are therefore needed to clarify underlying molecular mechanisms for low HDL-C levels due to LXR pathway inhibition.

We also demonstrated that cholesterol overload and hepatic LXR inhibition affected RCT. An HC diet resulted in enhanced fecal excretion of macrophage-derived cholesterol, mainly as neutral sterols rather than bile acids (Figure 1G-I). This might have been due to increased catabolism of HDL (Figure 2A & B) and an increase in size of the whole body cholesterol pool.^22^ Interestingly, hepatic LXR inhibition through treatment with Ad-Sult2b1 reduced fecal excretion of both macrophage- (Figure 1H) and HDL- (Figure 2E) derived cholesterol only under HC, suggesting that cholesterol efflux from peripheral tissues to HDL did not contribute to reduced RCT due to Sult2b1. Taken together with the observations in the experiments using DKO, hepatic Sult2b1 overexpression reduced HDL-C levels and RCT via LXR-dependent pathways. Moreover, attenuation of RCT due to hepatic LXR inhibition might have been attributable to reduced expression of the LXR-target genes ABCG5 and ABCG8. The fact that this phenomenon was observed only under cholesterol overload suggests that LXRs are important cholesterol sensors in the liver.

Hepatic ABCG5 and ABCG8 heterodimers reportedly localize on the apical side of hepatocytes ^23^ and play a crucial role in biliary excretion of neutral sterols, but not bile acids.^24^ Further, attenuated ABCG5 and ABCG8 expression due to hepatic LXR inhibition might result in reduced bile excretion of neutral sterols, which could be a main mechanism for inhibition of RCT in mice treated with Ad-Sult2b1, and DKO. This idea is supported by our finding that there was greater accumulation of macrophage and HDL-derived cholesterol in livers expressing Sult2b1 as compared to the control (Figure 1E and 2C). We also observed that Sult2b1 overexpression did not affect bile acid excretion into bile, indicating that changes in another LXR target gene, CYP7A1, contributed less to reduced RCT in mice treated with Ad-Sult2b1. In addition, as compared to the control, hepatic Sult2b1 overexpression increased macrophage- (Figure 1E) and HDL- (Figure 2C) derived cholesterol in the liver, but not in bile. This finding was inconsistent with that in DKO (Figure 4D & E) and although precise mechanisms remain unclear, systemic ABCG5 and ABCG8 deletion (especially in intestine) might be the reason for this difference.

Next, we observed that hepatic LXR inhibition increased apoB-containing lipoprotein levels, which was further enhanced under HC (Figure 1B & C). These findings were consistent with those in systemic LXRα knockout mice ^18^ and DKO.^11^ We also confirmed this phenomenon in DKO (Figure 4B), though it appeared to be less striking compared to that observed by Breevoort et al.^11^ In order to elucidate the underlying mechanisms, we performed an *in vitro* study using cultured hepatocytes treated with Ad-Sult2b1. This revealed that Sult2b1 overexpression resulted in increased production of apoB-containing lipoproteins (Figure 3). Although we did not perform a kinetic study using labeled apoB-containing lipoproteins, it is unlikely that Sult2b1 affects their catabolism because there was no significant difference in LDLR expression between Ad-Sult2b1 and the control (Figure 1A). Regarding regulation of intracellular cholesterol content, we observed reasonable changes in SREBP2 expression due to Sult2b1 overexpression. SREBP2 levels were reduced by both HC and Sult2b1 overexpression, indicating that these treatments increased hepatic cholesterol content. If so, LDLR expression should be reduced under such conditions in theory. These findings are supported by the study of Peet et al ^18^ who reported that LXRα deletion resulted in increased hepatic cholesterol levels and reduced SREBP2 expression but LDLR expression was unchanged. Taken together with the unreasonable changes in PCSK9 levels in mice treated with Ad-Sult2b1 that we observed (Figure 1A), this indicates that further studies are needed to clarify the molecular mechanisms for LXR-dependent (Figure 1B & C)/independent (Figure 5B) effects of Sult2b1 on apoB-containing lipoprotein levels.

In conclusion, liver-specific LXR inhibition by Sult2b1 overexpression in cholesterol-fed mice reduced circulating HDL-C levels and RCT in an LXR-dependent fashion. Our observations may therefore provide the basis for novel LXR targeted strategies against atherosclerotic diseases.

## Acknowledgements

We thank Drs Kaori Endo-Umeda and Makoto Makishima for kindly providing us with LXRα/β double knockout mice. We also appreciate Dr Takeshi Adachi for providing us with a fast protein liquid chromatography system.

## Sources of Funding

This study was supported by Foundation for Promotion of Defense Medicine.

## Disclosures

None.

## Highlights

- LXRs have been promising therapeutic targets; however, it remains unclear whether hepatic LXRs contribute to reverse cholesterol transport (RCT).
- Liver-specific LXR inhibition in mice was achieved by transduction with an adenoviral vector harboring Sult2b1.
- Hepatic Sult2b1 overexpression in cholesterol-fed mice reduced circulating HDL-C levels and RCT in a LXR-dependent fashion.
- These observations may provide the basis for novel LXR targeted strategies against atherosclerotic diseases.

## Nonstandard Abbreviations and Acronyms

ABCA1: ATP-binding cassette transporter A1
ABCG5: ATP-binding cassette transporter G5
ABCG8: ATP-binding cassette transporter G8
Ad-Luc: adenoviral vector harboring luciferase
Ad-Sult2b1: adenovirus expressing mouse Sult2b1
CEs: cholesteryl oleate
CYP7A1: cholesterol 7α hydroxylase
DKO: LXRα/β double knockout mice
FCR: fractional catabolic rates
HDL-C: high-density lipoprotein-cholesterol
LDL-C: low-density lipoprotein-cholesterol
LXR: liver X receptor
LSC: liquid scintillation counting
RCT: reverse cholesterol transport
Sult2b1: sulfotransferase family cytosolic 2B member 1
Wild-type: C57BL/6J mice

## Notes

### Competing Interest Statement

The authors have declared no competing interest.

